# Duplication accelerates the evolution of structural complexity in protein quaternary structure

**DOI:** 10.1101/2020.04.22.054783

**Authors:** Alexander S. Leonard, Sebastian E. Ahnert

## Abstract

Gene duplication, from single genes to whole genomes, has been observed in organisms across all taxa. Despite its prevalence, the evolutionary benefits of this mechanism are the subject of ongoing debate. Gene duplication can significantly alter the self-assembly of protein quaternary structures, impacting the dosage or interaction proclivity. Here we use a lattice model of self-assembly as a coarse-grained representation of protein complex assembly, and show that it can be used to examine potential evolutionary advantages of duplication. Duplication provides a unique mechanism for increasing the evolvability of protein complexes by enabling the transformation of symmetric homomeric interactions into heteromeric ones. This transformation is extensively observed in *in silico* evolutionary simulations of the lattice model, with duplication events significantly accelerating the rate at which structural complexity increases. These coarse-grained simulation results are corroborated with a large-scale analysis of complexes from the Protein Data Bank.

## Introduction

Self-assembly is a widespread phenomenon among proteins, with a substantial fraction of proteins assembling into multimeric complexes. These protein complexes exhibit a diverse range of topological configurations.

For the classification and analysis of protein complex topologies, it is often beneficial to simplify the representation of a complex to a weighted graph or a lattice model. Such coarse-grained representations have given rise to a comprehensive classification and enumeration of possible protein complex topologie*sAhnert et al. (2015)*; *Marsh et al. (2015)*. A lattice-based polyomino selfassembly model *Ahnert et al. (2010)* offers a qualitative resemblance to protein quaternary structure and a tractable genotype-phenotype map that nevertheless exhibits rich behaviour in evolutionary models *Johnston et al. (2011)*; *Greenbury et al. (2014)*. It should be pointed out however, that such models do not cover certain classes of proteins, such as intrinsically disordered proteins or those that require cooperative binding.

We focus on a recent extension to the polyomino model using binary strings as interfaces *Leonard and Ahnert (2019)*, which naturally introduces a coarse-grained approximation of interaction binding. Importantly, both symmetric homomeric and heteromeric interactions are possible, as well as combinations thereof, allowing for more general assembly dynamics than previously permitted in polyomino models. We additionally allow variable genetic length, with genotypes growing and shrinking dynamically through duplication and deletion respectively.

The importance of duplication in the structural evolution of proteins was unknown when first encountered in the 1970s, but was quickly realised to be ubiquitous *McLachlan (1979)*. Whether duplication is predominantly a driver of innovation *Magadum et al. (2013)*, distributor of subfunctions *Gibson and Goldberg (2009)*, safeguard against extinction *Crow and Wagner (2005)*, or merely a passive passenger *Kimura (1991)* remains more an open question. However, evolutionary histories are rife with duplication events, from single genes to whole genomes, and there is some consensus that its overall effects on evolution are therefore likely to be positive.

Less complex organisms, such as prokaryotes, may have evolved predominantly through other genetic mechanisms, e.g. horizontal gene transfer *Treangen and Rocha (2011)*. Eukaryotes, on the other hand, exhibit ample evidence of duplication-driven evolution *Lynch and Conery (2003)*. A prominent example is the kinetochore, which is a large, multi-protein structure in eukaryotes that is essential for cell division, and which formed through extensive gene duplications *Tromer et al. (2019)*. The homologous proteins that arise from gene duplication have been shown to vary considerably in terms of robustness against negative mutations and evolvability towards positive ones *Diss et al. (2017)*; *Dandage and Landry (2019)*.

Advances in genetic sequencing methods have made it straightforward to identify duplication events, but repeated duplication and mutation events mean that it is difficult to obtain a comprehensive overview of the evolutionary history of a genome. Tractable models of evolution are therefore invaluable for the study of duplication events, and produce predictions that can be verified using measurements of real protein structures.

### Polyomino self-assembly

In the polyomino self-assembly model, two-dimensional square tiles with binding sites on their edges self-assemble into complex shapes (polyominoes). Aside from the configuration of the interfaces on the tiles, the model also requires a specification of the interactions between the interfaces.

The original polyomino model *Ahnert et al. (2010)* used integer labels for binding sites, but has recently been generalised by representing the binding sites with binary strings of length *L Leonard and Ahnert (2019)*. A genotype can encode one or more subunits, where subunit binding sites are conventionally labelled in the clockwise direction.

A natural description for the interaction strength is related to the Hamming distance of two counter-aligned sites, such that adjacent bits that differ contribute to binding while those that match contribute nothing. The binding strength is thus the total number of differing bits, 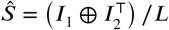, normalised by length such that ***Ŝ*** ∈ [0,1]. Two subunits bind to each other when this strength is equal to or greater than some threshold value ***Ŝ_c_***.

While this model is not designed to describe the thermodynamics of binding, the probability of an interaction occurring during assembly can be weighted by its strength. This results in stronger interactions binding more readily, as might also be expected in physical protein complexes. The dynamics of interest in this work do not depend on the precise relationship between interaction strength and binding probability. We therefore choose the simple relationship ***H* (*Ŝ_c_ – Ŝ*) · *Ŝ***, where ***H*** is the Heaviside step function with ***H*** (0) = 1, which offers analytical tractability.

These two ingredients are used to construct an *assembly graph*, which encodes all possible interactions between subunits for a given genotype. The assembly graph can be fed into the stochastic self-assembly algorithm as follows:

- seed structure with random subunit
- pick an “open” interaction on existing structure
- attach a subunit which binds to that interaction
- repeat previous 2 steps until no “open” interactions remain

The resulting lattice structure is a polyomino, taken here to be the phenotype that corresponds to the given genotype. For the purposes of our model there is an infinite number of copies of each subunit, and subunits bind irreversibly to the structure. Subunit composition within the polyomino is not considered in this work, and absolute translations or rotations of a polyomino are not considered distinct. Chiral counterparts of polyominoes are not considered distinct phenotypes, and as such these are *free* polyominoes. An example of an assembly is shown in *Figure 1*.

**Figure 1.**
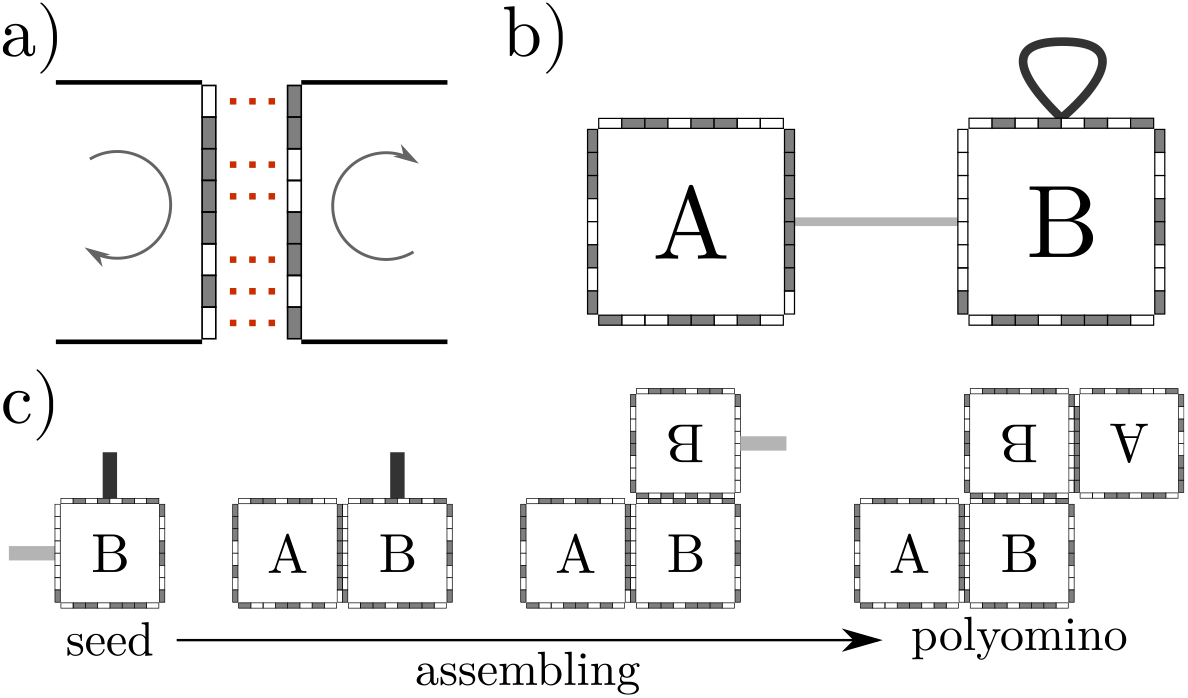
Self-assembly in the generalised polyomino model. a) A heteromeric interaction with red dots indicating contribution to binding strength between the oppositely directed binary strings, giving 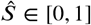. b) An assembly graph which contains one heteromeric (grey) and one symmetric homomeric (black) interaction. Subunit labels, interaction colours, and explicit binding sites are purely for clarity and not commonly shown for assembly graphs. c) A possible assembly pathway for the stochastic algorithm described in text, producing a tetramer.

Assemblies which eventually run out of “open” interactions and cease to grow are classed as *bound*, while those growing indefinitely are *unbound*. For two-dimensional lattice assembly, unboundedness is generally the result of having two or more distinct *cycles* in the assembly graph, which are repeating patterns of assembly. Assembly graphs can support complex cycles of different periodicity, where the simplest cycle is a self-interaction (***C***_2_ symmetry cycle).

Some assembly graphs routinely assemble the same polyomino, while others may stochastically yield different polyominoes upon repeated assembly. These are respectively termed *deterministic* and *nondeterministic* assemblies, where the phenotype can be defined as the most frequent polyomino. Both unbound and nondeterministic assemblies can be biologically unfavourable and are the underlying mechanism of a variety of proteopathic diseases.

### Interaction dynamics

Interactions can be grouped into two broad classes: symmetric homomeric, where a site binds to itself, and heteromeric, where two different sites bind. As symmetric homomeric interactions are assembly graph cycles, they substantially differ from heteromeric interactions in terms of genotypic evolvability and robustness. In addition, any mutation to a symmetric homomeric binding site changes the interaction strength by two increments, affecting both the location of the point mutation and the counter-aligned bit. This property of symmetric homomeric interactions is not unique to polyomino models, with greater variance in interaction energetics preferring symmetric homomeric formation extremely generally *Lukatsky et al. (2007)*; *Lukatsky and Shakhnovich (2008)*.

The simplicity of binary string binding sites allows the formation rates of both symmetric homomeric and heteromeric interactions to be calculated analytically. The formation of interactions is modelled by an absorbing Markov chain of interaction strengths, randomly walking below the strength threshold. Point mutations correspond to stepping to adjacent strength states. Symmetric homomeric interactions, which can only increment by two states, can be mapped to an equivalent chain of adjacent states by using 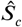 Extended details can be found in. This result has no closed-form solution, but an empirically derived approximation is given by:

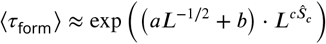

Where the coefficients should be estimated from the parameter window under investigation, with values of *a* = 0.71, *b* = 0.08, and *c* = 1.35 used in this work. Formation times grow exponentially with binary string length ***L*** and strength threshold ***Ŝ**_c_*. Symmetric homomeric interactions, with an effective interface length of 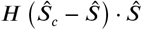, therefore take far fewer mutations on average to form.

Interaction loss can be understood using the same machinery, but on a reversed Markov chain with different initial conditions, also fully described in. Similarly to formation, symmetric homomeric interactions have double the variance and tend to be lost faster than heteromeric interactions. Loss, furthermore, takes place on incomparably shorter time scales to formation.

### Duplication-heteromerisation

Duplication can occur through multiple biological mechanisms, such as replication slippage or unequal crossing-over *Ohno (1970)*. Certain mechanisms may be more likely to copy entire chromosomes or just short sequences, but ultimately can still only duplicate existing genotypic segments. In this model, duplication copies an entire subunit, similar to gene-level duplication, including all its interactions. Immediately after duplication there are then two identical copies of a subunit.

The original and duplicated subunits can then evolve and accumulate mutations independently. Importantly, duplicating a subunit containing a symmetric homomeric interaction results in at least three interactions: two duplicate symmetric homomeric and one heteromeric interaction between the two copies. Only one of these interactions is required to maintain the phenotype, allowing the other interactions to neutrally drift, potentially out of existence.

There are numerous pathways of different surviving interaction configurations and their strengths, but not all are equally robust or evolvable. As shown in *Figure 2*, pathways ending with only a heteromeric interaction can most advantageously evolve a new symmetric homomeric interaction, and so have the highest “meta-fitness”.

**Figure 2.**
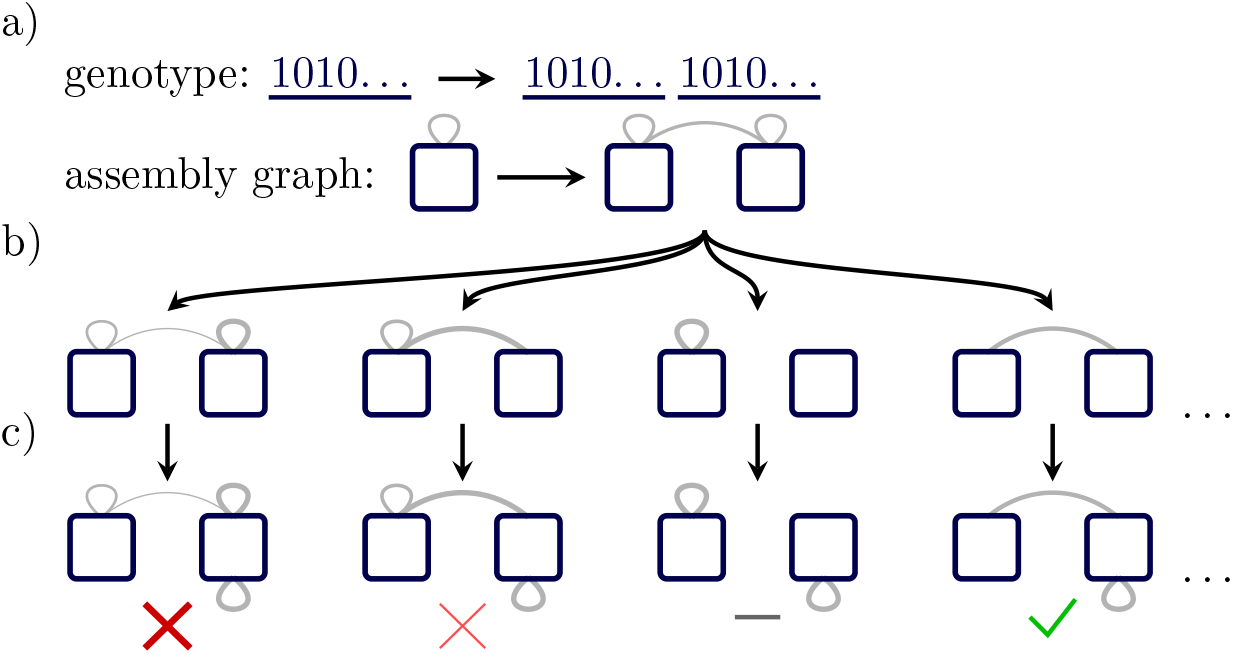
Duplication-heteromerisation pathway mechanism. a) Duplication events replicate regions of the genotype encoding a single subunit, duplicating any interactions involving that subunit. b) There are many configurations in which mutations can accumulate on the existing interactions, with interaction width indicating strength. All possible drifts which preserve at least one interaction are neutral. c) Forming a new interaction breaks the neutrality. From left to right, the fitness change is highly negative (unbound & strongly nondeterministic), slightly negative (weakly nondeterministic), neutral (same phenotype), and positive (bound & deterministic, larger phenotype).

### Complexity ceiling

Reasonably bound and deterministic assembly graphs can only support a limited number of interactions relative to the number of subunits *Tesoro et al. (2018)*. This limit can be recast in terms of the maximum “complexity” an assembly graph can support, as more interactions require more information to encode. Beyond this complexity, misassembly dominates.

This is particularly applicable to assembly graphs with cycles, as multiple cycles readily misassemble. Since the most likely evolutionary pathway starts with forming a symmetric homomeric interaction, a ***C***_2_ cycle, most genotypes then rapidly reach their complexity “ceiling”. At this point, a large portion of possible additional interactions will induce deleterious behaviour. In order for a structure to evolve further, this ceiling has to be overcome.

Duplication-heteromerisation allows a genotype to circumvent the complexity ceiling by transforming an assembly cycle into multiple single-use interactions. Such a genotype can now accept almost any new interaction, rather than being stuck at the bottleneck needing certain heteromeric interactions only. As a result, evolution can proceed at an accelerated rate when genotypes can duplicate.

## Results

Evolutionary histories were simulated to test the proposed pathway. Discrete generations of selection and mutation to a fixed-size haploid population formed the base of the evolutionary dynamics. Reproduction at each generation was asexual, with a fixed probability to flip each bit in the genotype and a separate probability to duplicate or delete a subunit. More fit individuals were selected proportionally more often to reproduce, with the fitness defined as the size of the most frequent polyomino weighted by how regularly it was assembled.

Duplication and deletion introduce dynamic genotype length and potentially rapid turn-over of subunits, and so tracking individual interactions overtime is no longer meaningful. Instead, “degenerate” interactions should be viewed as an ensemble, e.g. a symmetric homomeric interaction and a duplicated copy are fundamentally related, and should be considered together in order to capture the entire dynamics acting on their evolution. A simple approach to defining an ensemble is by the geometric position of the interactions on their subunits, with some examples shown in *Figure 3*.

**Figure 3.**
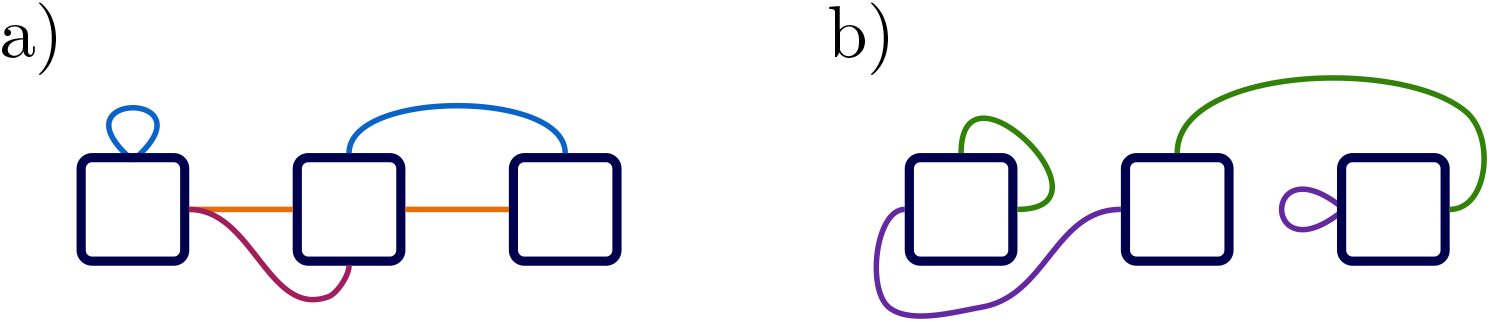
Grouping of degenerate interaction topologies. Two example assembly graphs with three and two interaction ensembles respectively in a) and b), where all interactions in an ensemble have the same colour. Two interactions can be degenerate even if one is intra-subunit and another is inter-subunit, due to the duplication-heteromerisation process. Any interaction properties, such as interaction strength or sequence similarity, should be considered over an entire degenerate ensemble, e.g. all purple-coloured interactions.

Heteromeric interactions can be further classified into homologous, where binding sites share ancestry, and non-homologous, where they do not. Such shared history can only occur through subunit duplication. Non-homologous heteromeric interactions are formed *de novo*, with no existing relationship between the involved subunits. Understanding the evolution and frequency of these different interaction types plays a crucial role in uncovering the purpose and impact of duplication.

### Interaction homology

Sequence similarity is a critical tool in identifying protein homology, measuring how many amino acids match or nearly match between two protein sequences. In the context of binary strings, interaction sequence similarity measures how identical two binding sites are through the Hamming distance, as the bits either match or are opposite. Sequence similarity of arbitrary binary strings can be predicted with a binomial distribution, ***B* (*L*,.5)**.

Symmetric homomeric interactions always have maximum sequence similarity, by definition, as they interact with themselves. Heteromeric interactions do not have such a constraint, and can take any possible value. Two unrelated sequences are expected to have similarity near the mean of the binomial distribution, 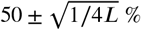. However, finding a heteromeric interaction with high similarity (» 50%) suggests they are not two arbitrary binding sites, but instead share evolutionary origin. Example simulated evolutions of interaction similarity are shown in *Figure 4*.

**Figure 4.**
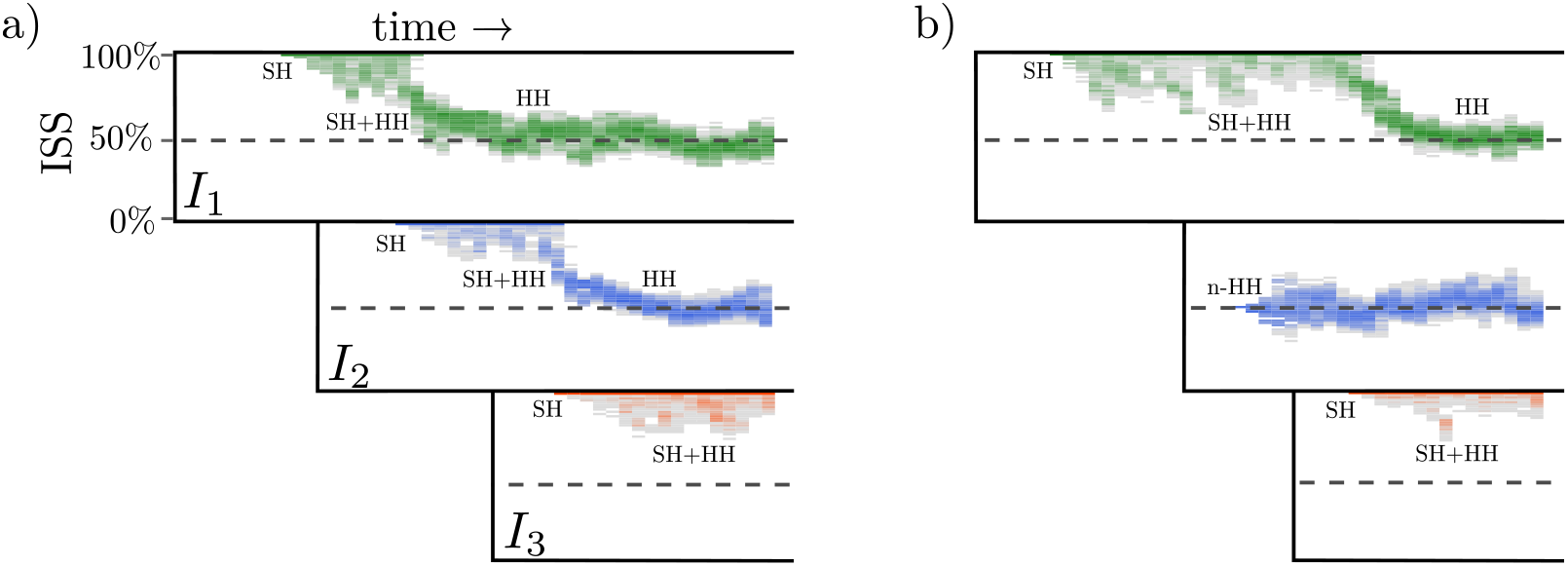
Interaction sequence similarity evolutions. Interaction sequence similarity (ISS) can change during evolution, shown here for three sequentially formed interactions (*I*_1_,*I*_2_, and *I*_3_) over two independent simulations in a) and b). Both simulations initially form a symmetric homomeric interaction (SH), which later coexists with its duplicate-drift homologous heteromeric interaction (HH). Forming a new symmetric homomeric interaction fixes the purely homologous heteromeric state. Note in b), forming a non-homologous heteromeric interaction (n-HH) as *I*_2_ does not break the neutral symmetry of the SH+HH state, which continues to duplicate and drift.

Duplication probabalistically occurs at all times, but is most noticeable after a symmetric homomeric interaction forms. Immediately after duplicating a subunit containing a symmetric homomeric interaction, heteromeric copies appear. As these heteromeric copies accumulate mutations and neutrally drift, their sequence similarity correspondingly drifts towards heteromeric expectations of 50%. These interactions can also be lost by drifting below the strength threshold, provided at least one interaction copy survives. This pattern of drift results in “smears” in sequence similarity, as heteromeric interaction differentiate themselves but then are lost to interaction or population dynamics.

Only when a new symmetric homomeric interaction forms is that neutral symmetry broken. At this stage, the heteromeric copy of an originally symmetric homomeric interaction quickly fixates in the population and fluctuates around the expected mean. The recently formed symmetric homomeric interaction then undergoes smearing drift, and the process can repeat. Note that forming an non-homologous heteromeric interaction (*Figure 4b*, *I*_2_) has minimal impact on symmetric homomeric interactions which continue smearing drift.

### Interaction Composition

Evolutionary simulations are highly stochastic, and the identification of single events is informative but insufficient to understand the diversity of dynamics at play. Averaging over multiple independent simulation provides a more comprehensive picture. Interaction types, averaged over many assembly graphs, can easily be captured at every phenotype transition step, providing time-resolved tracking of composition.

Without duplication, interactions are static and have no ability to change types, while with duplication symmetric homomeric interactions can heteromerise. Although possible at any time, duplication only meaningfully influences subunits with existing interactions. Therefore, any consequences of duplication are only expected to arise after an interaction has formed. Interaction compositions reflect these predictions exactly, seen in *Figure 5*.

**Figure 5.**
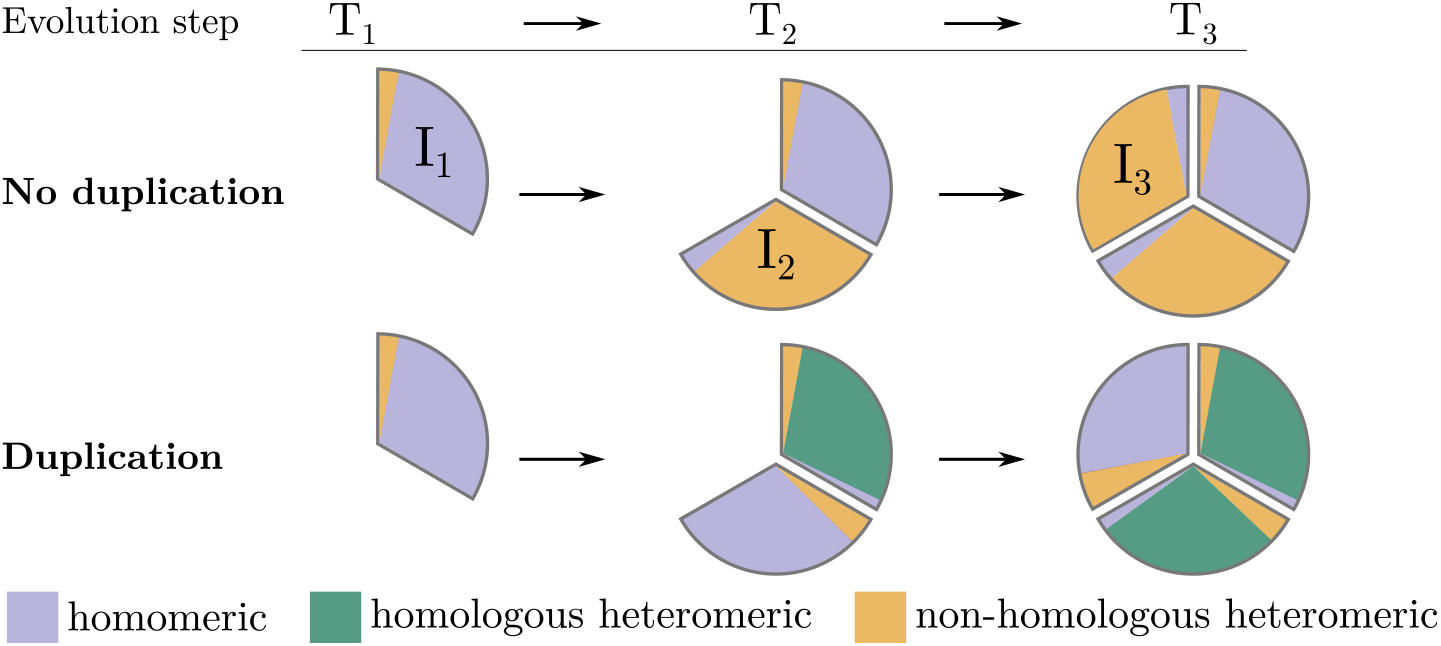
Assembly graph interaction composition over time. Assembly graph composition was recorded at each phenotype transition for the first three interaction (*I*_1_,*I*_2_, and *I*_3_) formed at three sequential times (*T*_1_, *T*_2_, *T*_3_ respectively), averaged over 1500 simulations. Without duplication, interaction types are static and do not change throughout evolution; later periods of evolution are predominantly driven by non-homologous heteromeric interactions. When duplication is allowed, symmetric homomeric interactions can transition to homologous heteromeric to facilitate evolvability, as strongly observed for the *I*_1_ interaction at time *T*_2_ and likewise *I*_2_ at *T*_3_.

After evolving through three phenotypes, the average assembly graph is comprised of about one symmetric homomeric and two heteromeric interactions, regardless of duplication. When duplication is allowed, however, the heteromeric interactions are overwhelmingly homologous. There is a diminishing returns effect in duplication due to dosage (see Discussion) as it becomes more difficult to duplicate and heteromerise a larger complex, but the effect is small for only a few interactions.

### Accelerating evolution of complexity

Tracking the number of interactions over time offers a reliable proxy to measuring the evolution of structural complexity, *C*. Again due to the stochasticity inherent in single evolutions, averaging over many simulations provides a more stable trend that is suitable for interpretation. Although the Markov chain predictions for interaction formation have enormous variance, at times comparable to the mean, these expressions yield approximate checkpoints for the complexity quantity. The quicker symmetric homomeric interaction formation time, 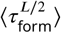 should coincide with 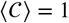. The second interaction to form should be heteromeric, and so 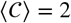 is expected around 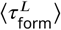.

As mentioned previously, duplication only realistically impacts evolution after the formation of the first interaction, and so the initial rates of evolving complexity should be comparable. Once duplication-heteromerisation comes into play, however, evolution should be accelerated with 〈c〉 growing faster than expected. That means for a given period of evolution, simulations with duplication allowed should be at comparatively higher complexity. This is precisely observed in *Figure 6*.

Evolutionary runs were restricted to a maximum of three interactions due to finite simulation resources, which is why the complexity asymptotically levels off at long time scales. As both sets of simulations converge to the imposed limit, the complexity gap diminishes to zero. However, smaller samples corroborate that the gap continuously widens when no constraint is placed, implying that duplication pathways repeatedly outpace standard evolution. Furthermore, comparable simulations using insertion of random subunits rather than duplication do not display any such acceleration. As such, the benefit appears localised to duplication-heteromerisation pathways rather than additional genetic length.

**Figure 6.**
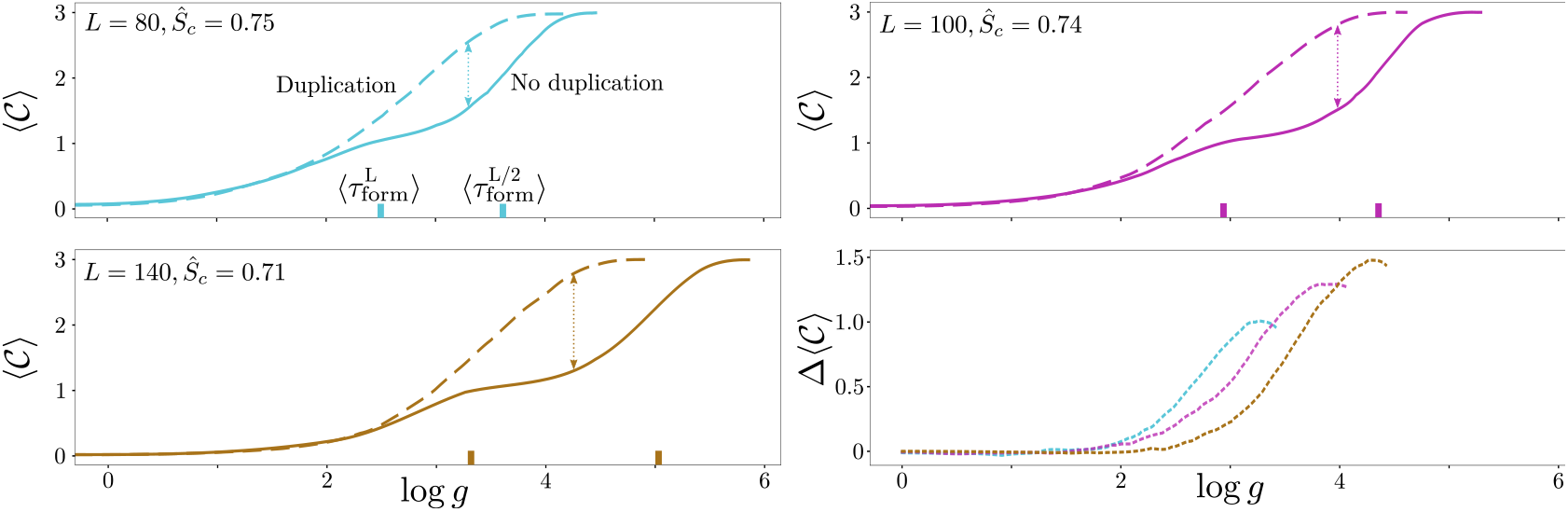
Evolution of structural complexity. Structural complexity, *C*, can be approximated by the number of interactions in an assembly graph. Trends are stable after averaging over 1500 simulations, up to a computationally imposed limit of three interactions. Note that a visually parallel line but at larger log *g* corresponds to a smaller slope due to logarithmically scaled generations. Duplication accelerates growth of complexity, measured by the gap between evolution ability with and without duplication, Δ〈*c*〉. As the duplication-heteromerisation becomes increasingly advantageous, evolution gains a greater acceleration.

Importantly, as can be seen in the bottom right panel of *Figure 6*, the observed evolutionary advantage increases with binding site length ***L***. This arises from the scaling “size” of the heteromeric bottleneck, where the ratio of times to form a symmetric homomeric and heteromeric interaction diverges quasi-exponentially, given the approximate form of 〈*τ*_form_〉. The evolutionary advantage of a parameterisation can be predicted *a priori* through the above ratio and the fraction of duplication events that neutrally drift to the heteromeric state. Extended details on these calculations and demonstration of the universality of the pathway are given in.

### Protein data bank analysis

The lattice self-assembly model is a limited representation of real protein complexes. However, the simulations detailed above generate several qualitative hypotheses that can be compared to complexes taken from the Protein Data Bank (PDB) *wwPDB consortium (2018)*. This data can be used to corroborate the hypotheses, even without access to the specific evolutionary histories of individual proteins.

We can identify homologous and non-homologous heteromeric interactions by comparing the domains assigned to each protein subunit. Domains hierarchically cluster protein subunits according to structural similarities. They typically correspond to compact units of folding which are conserved in evolutionary histories. CATH *Dawson et al. (2016)* is a major domain database, which assigns (potentially multiple) four segment codes to protein subunits. If two proteins share domains, they are in the same homologous superfamily and broadly overlap in terms of their evolutionary ancestry.

The set of proteins analysed here is derived from a previously curated dataset *Ahnert et al. (2015)*. Proteins that had missing or incomplete bioinformatic data, CATH domains or macromolecular interfaces, are removed, with representants taken at 90% subunit sequence similarity. Full details on compiling the protein complexes from the dataset are provided in the Methods section.

### Domain co-occurrence statistics

One of the simplest observations is the excessive correlation of domains across heteromeric interactions, i.e. the overabundance of homologous heteromers. A permutation test, randomly shuffling the domain assignments on each subunit and recalculating the quantity of homologous and non-homologous heteromeric interactions, is shown in Table 1. Fisher’s exact test yields an overwhelmingly significant confidence, *p* ~ 5 × 10^-235^, that subunits involved in heteromeric interactions are more commonly homologous than naively expected. Additionally, only about 4% of domains in homologous heteromeric interactions did not also exist in a symmetric homomeric interaction, while it was double that for the non-homologous case.

**Table 1.** Permutation test on heteromeric domains. CATH domain overlap for 3658 heteromeric interactions, with any or no domain overlap. Interactions were classified with domains taken from the data as well as randomly shuffled from the pool of 7316 heteromeric subunits.

|  |  | Domains |  |
| --- | --- | --- | --- |
|  |  | Homologous | Non-homologous |
| Pairing | CATH | 1528 | 2130 |
|  | Shuffled | 342 | 3316 |

In addition, many protein complexes are comprised of a mixture of interaction types. Approximately one quarter of complexes containing at least one heteromeric interaction also had a least one symmetric homomeric interaction. Out of 594 complexes with multiple heteromeric interactions, 73% of them contained a homologous heteromeric interaction. The average composition for these 594 proteins was about 2.0 homologous and 1.5 non-homologous heteromeric interaction, with *p* = 6.6 × 10^-12^ that any complex has a greater likelihood to have more homologous than non-homologous interactions. Such results qualitatively agree with the model simulations, showing that homologous heteromers are a significant contributor to the evolution of larger complexes.

### Interaction buried surface area

One clear trend from model simulations is that longer binding site lengths ***L*** and higher strength threshold ***Ŝ**_c_* result in a higher proportion of homologous heteromeric interactions. While the physical binding of protein subunits has complex dependencies on kinematics and interaction energies of their buried surface area (BSA), the two model parameters ***L*** and ***Ŝ**_c_* controlling interactions are reasonable as a first order approximation. BSA is influenced by many additional factors, such as steric exclusion or cooperative binding, but correlates strongly with binding affinitsy *Chen et al. (2013)*.

Without the ability to rerun evolution, the protein data cannot show that larger protein binding sites more frequently evolved via homologous heteromeric interactions. However, assessing the homology and BSA of heteromeric interactions taken from the protein data allows the converse question to be asked: are larger BSA heteromeric interactions more commonly homologous or non-homologous? Although the arguments seem similar, this question does not isolate the influence of BSA size to interaction binding, and results should be interpreted cautiously. Nonetheless, the gathered results clearly display a statistically larger BSA (“α ***L***”) for homologous compared to non-homologous heteromers seen in *Figure 7*, supporting a key prediction from the model.

The data contains many homologous heteromeric interactions related to immunoglobulin proteins with high BSA, present in the upper peak. This immunoglobulin domain (CATH: **2.60.40.10**) is the most common recorded domain, both in the dataset as well as in CATH. Statistical significance is still retained after removing any protein subunits with immunoglobulin domains (*p* ≈ 7 × 10^-10^, CLES=.59), filtering at a 70% cluster similarity threshold (*p* ≈ 5 × 10^-9^, CLES=.57), as well as combinations of these filters.

### Subunit binding sequence alignment

Alongside the structural and atomic coordinate data from the PDB and interaction homology from CATH, residue-level details from a database of crystallographic interactions (PDBePISA *Krissinel and Henrick (2007)*) are critical to substantiating the duplication-heteromerisation pathway. Examin-ing the specific residues involved in heteromeric interactions, where two subunits may have an excessive similarity, yields more decisive proof that the interaction was once symmetric homomeric and has since duplicated and heteromerised. The subunit binding sequence alignment (SuBSeA) analysis combines optimal sequence alignment with associating individual residues with the binding subunits. Full details on this procedure can be found in the Methods section, with a diagrammatic overview in *Figure 8*.

**Figure 7.**
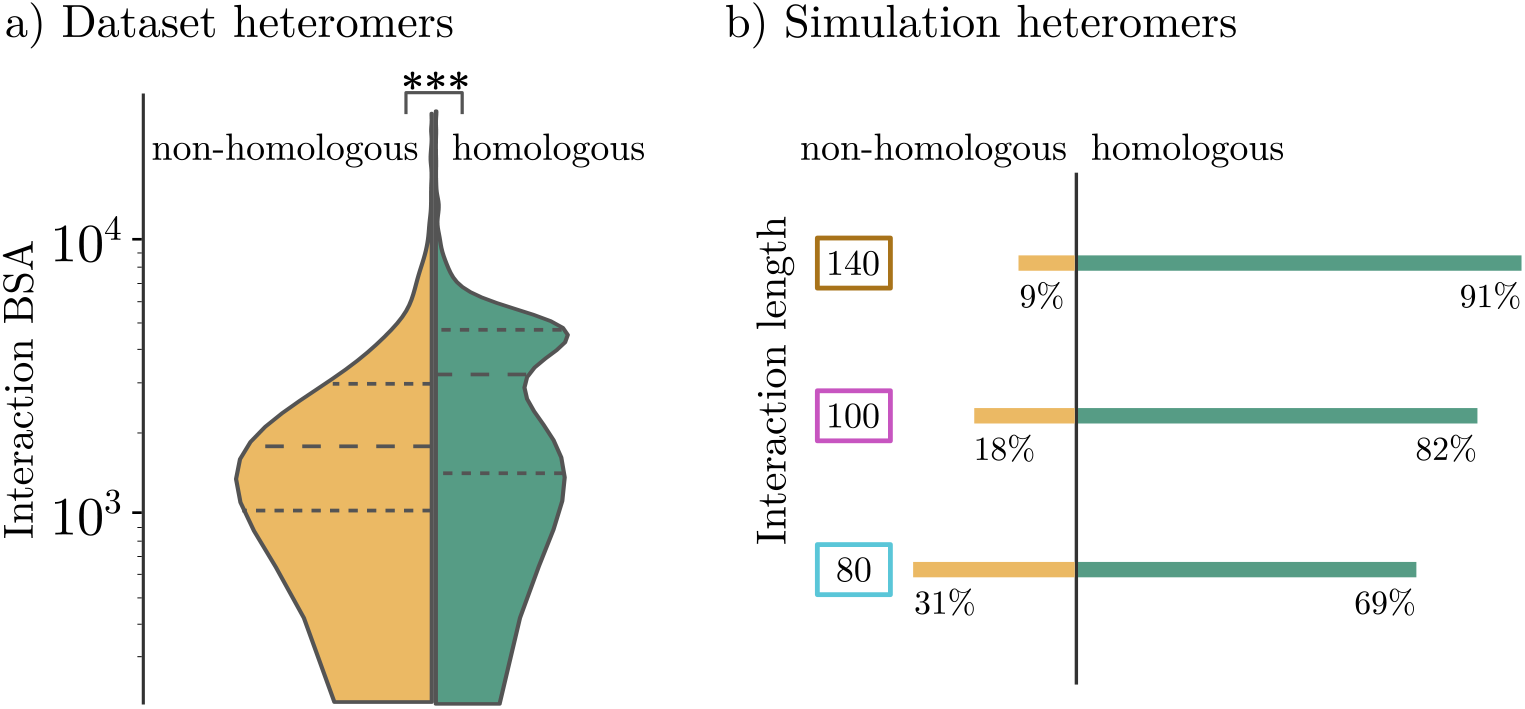
Binding surface area distributions. a) Homologous heteromeric protein interactions have statistically larger buried surface area than non-homologous interactions according to a Brunner-Munzel test: *p* ~ 10^-47^. The common language effect size (CLES) is .64, i.e. the homologous BSA will be larger in 64 out of 100 random pairs. b) Heteromeric interactions from *in silico* simulations skew towards homologous ancestry as ***L*** increases, parallel to the dataset results. Non-homologous heteromeric interactions can still form at long interaction lengths, but quickly become rare.

**Figure 8.**
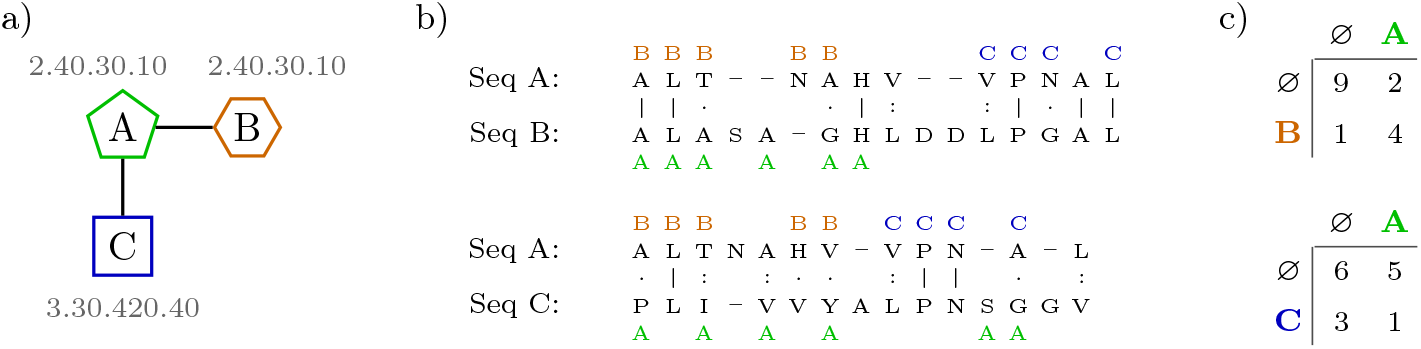
Subunit Binding Sequence Alignment comparison method. a) A trimeric protein where subunits A and B have a homologous heteromeric interaction while A and C have a non-homologous one, according to their example CATH domains in grey. b) After sequences are aligned, residues are clearly marked if they are involved in an interaction with another subunit. Identical, similar, and unrelated residues are indicated by ‘|’, ‘:’ and ‘.’ respectively. c) Aligned binding residue partners are counted, e.g. how many times a ‘C’-interacting residue is opposite an ‘A’-interacting residue, or a ‘B’-interacting residue opposite a non-interacting residue (denoted ‘ø’). The results are stored in a matrix **M**, which can then be used to infer duplication-heteromerisation confidence.

Confidences are assigned for each heteromeric interaction, which are grouped into homologous or non-homologous if they do or do not share domains, as previously. Survival functions for the distributions of these two groups are shown in *Figure 9*. These functions measures how much of a population is above some confidence level, and are related to cumulative distribution functions as SF = 1 – CDF. Crucially, homologous heteromeric interactions display substantially stronger confidences, giving credence to ancestral symmetric homomeric interactions which explicitly underwent duplication-heteromerisation.

Fewer homologous heteromeric interactions failed to find any alignment, which results in log *p*\* = 0, compared to non-homologous interactions, 42% and 75% respectively. In addition, the power law approximation shows the SuBSeA confidence in homologous heteromeric interactions decays roughly twice as slowly as the non-homologous case. Nearly a third of the homologous alignments were more significant than the common threshold of *p*\* = 0.05 (-log *p*\* ≈ 1.3), compared to only about 10% for non-homologous interactions.

**Figure 9.**
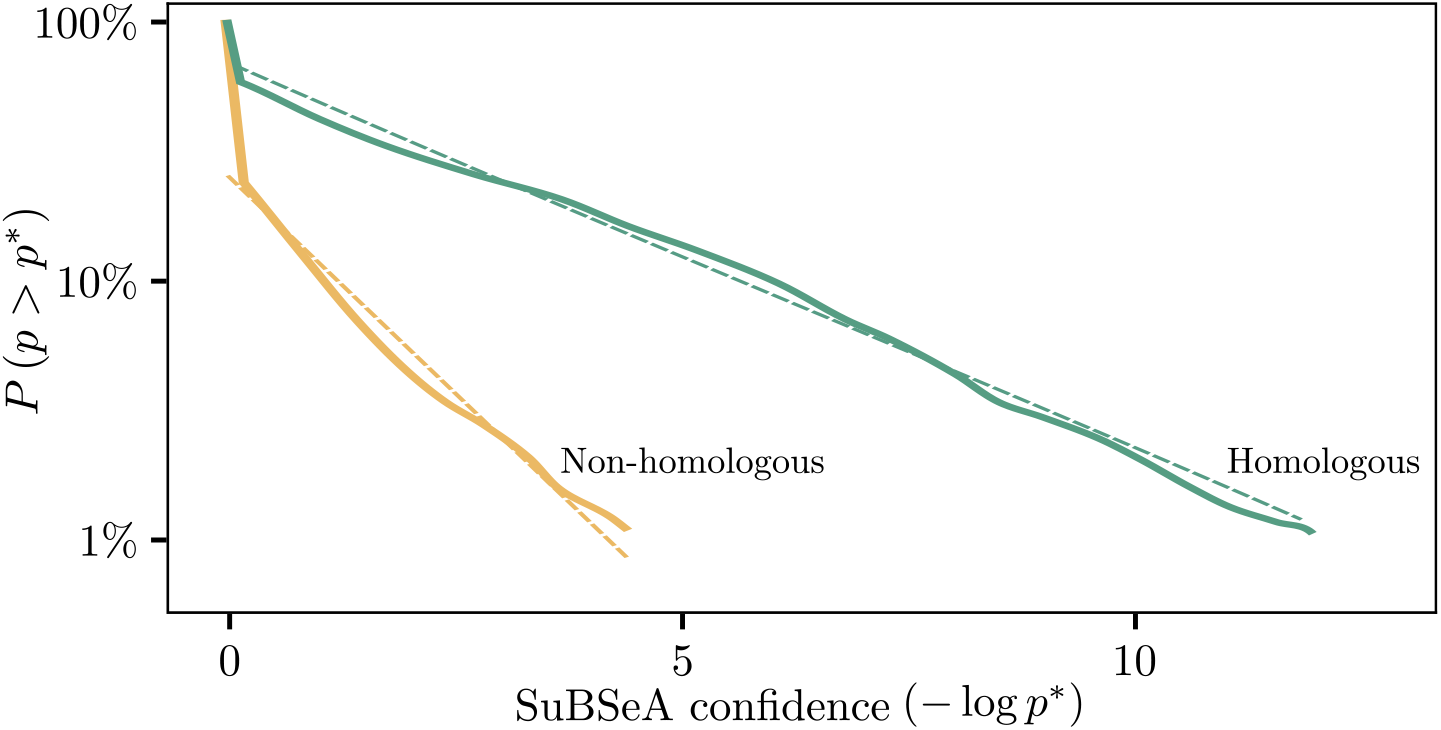
SuBSeA confidences for heteromeric interactions. SuBSeA confidences calculated for 1528 homologous and 2130 non-homologous heteromeric interactions from the dataset. Homologous interactions displayed stronger confidences, evidenced by the slower decaying survival function, i.e. more homologous interactions had confidences greater than any arbitrary *p*\* compared to non-homologous interactions. The survival functions can be approximated with a power law for – log *p*\* > 0 using the form « *p*\*^-*ξ*^, with exponents of 0.15 and 0.34 respectively.

Homologous heteromeric interactions also displayed stronger SuBSeA confidences when compared against domain-matched symmetric homomeric interactions, seen in S1 Fig. Identifying symmetric homomeric precursors provides additional confirmation that homologous heteromeric interactions specifically originated through duplication-heteromerisation.

## Discussion

While this lattice polyomino self-assembly model with binary binding sites serves as a reasonable coarse-grained representation for a broad range of well defined proteins, it is a highly idealised model. However, the duplication pathway proposed here is non-specific, based on general imbalance between symmetric homomeric and heteromeric binding properties *Lukatsky et al. (2007)*; *Lukatsky and Shakhnovich (2008)* and an assumption that duplication events are (approximately) neutral.

Other dynamics associated with duplication were inherently present in our simulations, such as protein dosage which can have a role in evolution outcomes *Edger and Pires (2009)*. Duplication of a subunit in our model immediately increases the likelihood that a nondeterministic assembly will integrate that subunit, as the number of relevant interactions has doubled. This generally only has a minor influence on fitness weighting, with a slight purifying pressure. More importantly, dosage introduces a form of self-regulation to constrain the genotypic length, where excessive copies of any given subunit makes the genotype so robust to negative mutations, that it struggles to evolve from positive ones.

Existing studies have reported on paralogous protein complexes arising from duplicating symmetric homomeric interactions. Pereira-Leal et. al focused on the existence of modularity in protein-protein interaction networks *Pereira-Leal et al. (2007)*, while Marchant et. al explored the role of heteromerisation in functional divergence *Marchant et al. (2019)*. Our results offer another perspective to the same observations: duplication-heteromerisation enables additional, faster pathways of evolving further interactions. This process involves distributing a symmetric homomericinteraction across two duplicated interfaces, each individually losing their self-interactions. As such, this pathway fits nicely under the concept of duplication-degeneration-complementation *Force etal. (1999)*.

This pathway is also strongly stochastic, randomly depending on the duplicated interactions accumulating mutations. Often in the simulations, the homologous heteromeric interaction was lost or a deleterious phenotype evolved, such that duplication only provides the pathways rather than imbuing them with a selective advantage. This likewise matches observations in the literature, where the fate of duplication is influenced by local contexts but largely succumb to random mutations *Teufel et al. (2018)*; *Lynch and Conery (2000)*.

Crucially, several interactions in the non-homologous distribution tail of *Figure 9* highlight the uncertainty in domain assignments. For example, the largest outlier (PDBid: 3F3P) has single digit sequence similarity between subunits J and L, but substantial structural similarity around the interacting region and a SuBSeA confidence of *p* ~ 10^-23^. These subunits may have duplicated and diverged too far in the past to be assigned homologous domains, with subunit J and L having “propeller” and “sheet” domain architectures respectively.

Symmetric homomeric interactions are an incredibly widespread type of interaction *Marianayagam et al. (2004)*, and have been the sole focus of the duplication-heteromerisation pathway thus far. These interactions impart the protein complex with ***C***_2_ symmetry; however, the concepts outlined here apply far more generally. Duplication-heteromerisation can act generally on any cyclic assembly to reduce the inherent symmetry through subfunctionalisation of the involved interactions. A homotetramer complex (***C***_4_) can duplicate-heteromerise to a homodimer of heterodimers (loosely 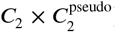), where the heterodimer is comprised of homologous subunits. This complex can transition even further to an asymmetric complex of four homologous heteromeric subunits.

Such behaviour can conceptually be extended to three dimensions and other symmetries, e.g. dihedral groups. As the duplication-heteromerisation pathway relies on neutral drift, these more complex cycles and symmetries almost certainly reach the evolvable heteromeric state less frequently. However, this is just a rescaling factor of the time spent in the drifting phase rather than any mechanistic difference.

Thousands of likely candidates are easily identified through browsing the PDB by pseudosymmetries. One example case is the complex PDBid: 3WJM, with two unique subunits forming a heterohexamer with ***D***_3_ pseudosymmetry. Both unique subunits share the same two CATH domains and have just below 80% sequence similarity. These subunits have SuBSeA confidences between the different subunit interaction conformations of 4 × 10^-7^ to 2 × 10^-34^. As such, this “dimer of trimers” probably evolved through cyclic interactions which have since duplicated and diverged into heteromeric states. This “X of Y” composition of complexes has been widely observed *Pereira-Leal et al. (2007)*; *Marsh and Teichmann (2014)*, and the proposed pathway is entirely complementary to these existing studies.

Another example is the complex PDBid: 2IX2, which forms a heterotrimer from three unique subunits but with ***C***_3_ pseudosymmetry. The three subunits average around 50% sequence similarity between any two. This does imply homology, typically taken above 30% similarity, but is relatively weak evidence *Pearson (2013)*. Comparing the interactions directly gives SuBSeA confidences of around 10^-7^, providing stronger evidence of shared ancestry. Furthermore, this is an example of an asymmetric homomeric interaction and a symmetry that is not possible in square geometry. These protein complexes and the relevant heteromeric interactions are shown in S2 Fig.

Genetic duplication has been involved in evolution across all domains of life, but is especially pervasive in more complex eukarya. There are various explanations on why duplication is so widespread, with many being complementary and possibily all part of the larger truth. However, without being able to replay the evolution of life, it is challenging to examine these reasons in depth. Here we have examined duplication in the context of protein self-assembly, using an abstract but powerful lattice polyomino model.

Duplication-heteromerisation offers a repeatable pathway to rapidly evolving structural complexity, by transforming fast-forming symmetric homomeric interactions into evolvable heteromeric interactions. Evolution of *in silico* self-assembling polyominoes made frequent use of this duplication pathway. Similar markers of duplication-heteromerisation were observed in a broad analysis of protein complexes from the PDB, finding conclusive evidence of binding residue alignment in homologous heteromeric interactions.

The problem of why duplication is seemingly so important to be so common remains unresolved, but our proposal for a simple but efficient evolutionary pathway offers a supplementary perspective that coexists with previous literature and observations.

## Methods

### Simulations

A population of haploids asexually reproduce at discrete generations, with a fixed population size of 100 individuals to minimize fluctuations but retain computational efficiency. Each bit in the genotype had a fixed independent probability to flip, chosen as *μ* = 1/(4*L*). Duplication and deletion were balanced such that there was no expected net growth unless driven by a fitness advantage. Multiple duplication and deletion rates were tested without any qualitative difference, and the final value of 0.05 per subunit per generation was chosen for optimal efficiency.

Each new generation was stochastically selected from the previous generation based on a fitness-proportional roulette wheel method. More fit individuals were proportionally more likely to reproduce and so had a better chance to eventually fixate. Fitness was determined as the weighted sum of each assembled polyomino size. For example, assembling a tetramer with frequency 0.75 and a dimer with frequency of 0.25 resulted in a fitness of *f* = 0.75·4+0.25·2 = 3.5. Most simulations evolved for 500,000 generations, except ***L*** = 60, ***Ŝ_c_*** = 0.83 and ***L*** = 140, ***Ŝ_c_*** = .71 which evolved for 1,000,000. This was due to the longer formation expectation times, such that each simulation on average would evolve three interactions.

### Phylogeny

Degenerate topologies were grouped by the geometric position of the interactions on subunits. For example, any interactions which occur between the top of a subunit and the top of a subunit are grouped together, or the left of one and the bottom of another. This grouping strategy eliminates the vagueness of when duplicated interactions have diverged enough to be differentiable, or if an interaction drifts below and then back above the strength threshold which interrupts continuous tracking. Since simulations are constrained to evolving a small number of interactions, the chance that two independent interactions would evolve with the same geometric configuration is low and is an insignificant source of noise compared to the signal.

### PDB dataset

The set of proteins analysed here is derived from a previously curated dataset *Ahnert et al. (2015)*. All entries with at least one symmetric homomeric or heteromeric interaction are included from the original and extended sets of bijective protein complexes. Only complexes that had data entries for FASTA amino acid sequence, PDB atomic coordinates, CATH domains, and PDBePISA macromolecular interfaces are kept. Redundant proteins interactions are filtered out using precompiled BLASTClust clusters (90% subunit sequence similarity, unless stated otherwise) available from the PDB. Only one representant interaction per combination of cluster IDs is allowed, picked randomly if multiple exist, to mitigate any abundance bias of proteins recorded in the PDB.

Similarly, protein complexes which contain many repetitious interactions, conceptually cycles in an assembly graph, have all interactions between identical subunits relabelled into a single interaction. If interaction sizes were slightly different between subunits, e.g. impacted by different conformations etc., an average value is taken. While there are many inherent biases and flaws that cannot be excluded within the databases, curated dataset, and data scrubbing methods, any overall trends present are reasonably robust enough to instil confidence in any conclusions.

### SuBSeA analysis

The SuBSeA analysis combines two datastreams: sequence alignment and macromolecular interfaces. Sequence alignment is straightforward, using the Needleman-Wunsch algorithm as implemented by EMBOSS needle *Madeira etal. (2019)* with default parameters. Binding residues are taken from the macromolecular interfaces from PDBePISA and converted to an intermediate format to track binding residues by subunit. The two datastreams are then overlaid, such that binding residues are aligned according to optimal sequence matching.

Co-occurrences of binding residues are then tracked in a matrix **M**, where each element is the number of aligned residues between two given subunits. In the common situation where one residue is not aligned opposite another target-interacting residue, it is tracked in a catch-all null column/row, ø. The co-occurrence counts and the marginal distributions approximate the probability that the alignment of an interaction could happen by chance. The probability for an element in the matrix is given by:

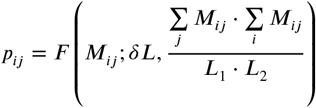

where ***F*** is the cumulative binomial distribution, ***L***_1_ and ***L***_2_ are the sequence lengths of the first and second subunits, and ***δL*** is the length of the overlapping alignment region.

If only null column and rows are present, the interaction is trivially assigned a unit p-value. Otherwise, the lowest probability (inferring the strongest likelihood of two interfaces being related) is stored and the relevant row and column are removed from the matrix. This process continues iteratively until there are no surviving rows or columns to sustain the matrix. The series of stored p-values is then combined using the Fisher method, although other methods can be used, e.g. Stouffer’s or Tippett’s, to determine the overall probability of interaction homology between two heteromerically linked subunits.

---

**Figure S1.**
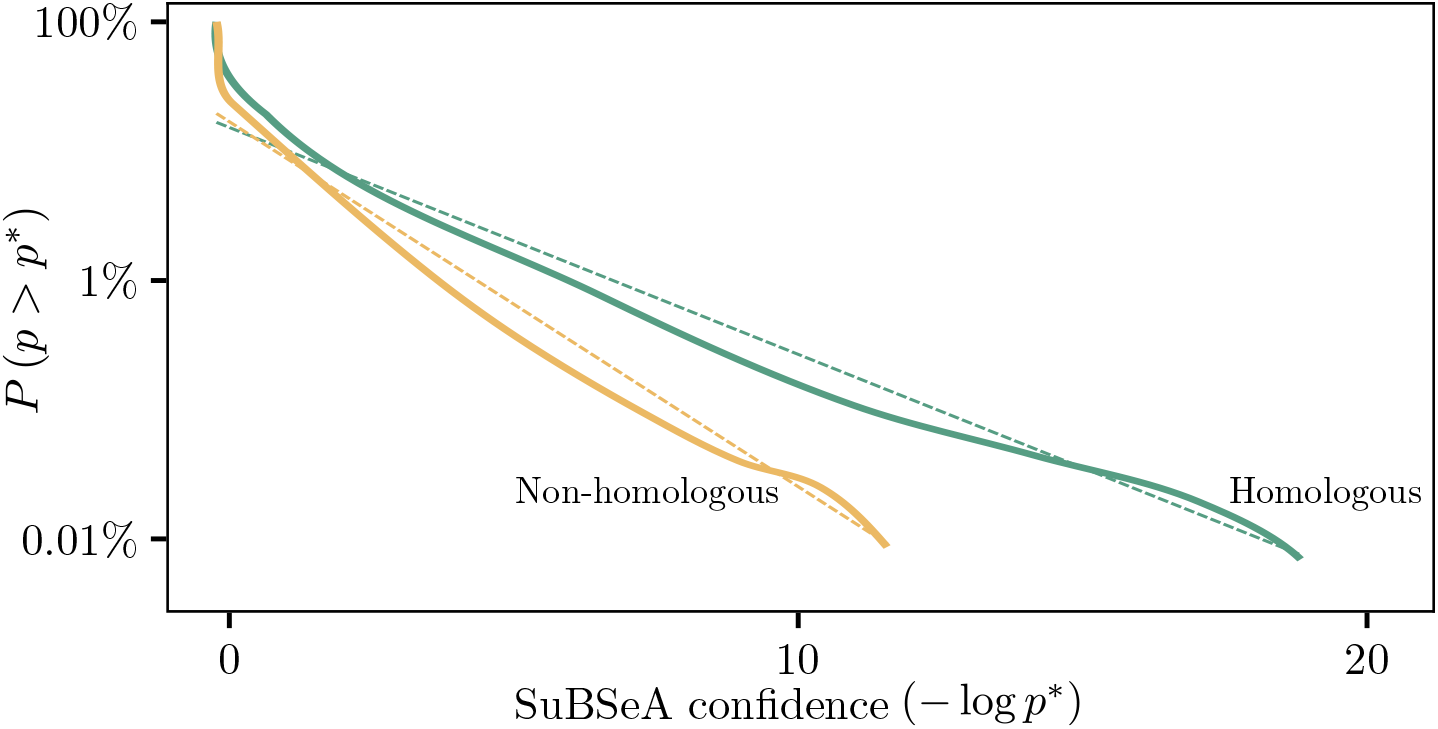
SuBSeA alignment for heteromeric interactions and predicted symmetric homomeric progenitors. Subunits with heteromeric interactions can be compared against symmetric homomeric complexes which share CATH domains. As in *Figure 9*, binding alignments which are homologous are statistically more confident. Since there are many more symmetric homomeric-heteromeric comparisons possible, the survival function is stable down to a lower percentage and higher *p*\*. The percentages of interactions failing to find any alignment were nearly identical to *Figure 9*, as 46% and 73% for homologous and non-homologous respectively (cf. 42% and 75%). The power law fits are approximately 0.22 and 0.33 respectively, (cf. 0.15 and 0.34).

**Figure S2.**
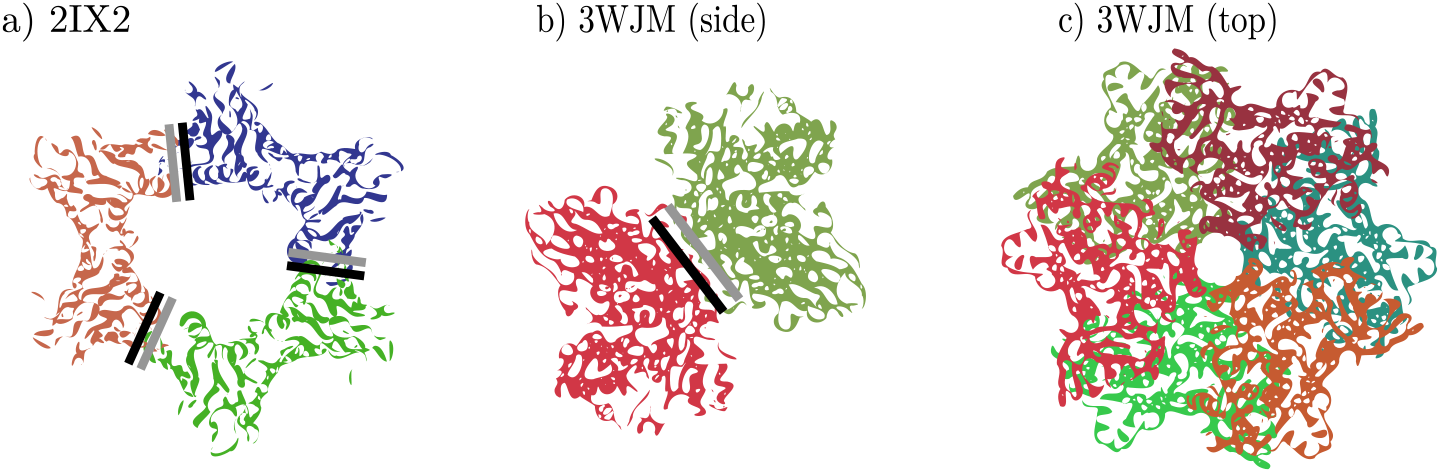
Duplication-heteromerisation in generalised geometries and symmetries. Two example protein complexes found through the PDB’s pseudosymmetry browser, where the binding alignments are highlighted with black and grey bars. a) PDBid: 2IX2 has ***C***_3_ symmetry, with heteromeric interactions between three distinct subunits. However, sequence similarity and SuBSeA confidences suggest this was originally an asymmetric homomeric interaction. b) A side-view of PDBid: 3WJM shows the dimeric heteromeric interaction present in this complex. The heteromeric interaction forming the dimer again has strong SuBSeA confidences, implying it was once symmetric homomeric. c) The top view of PDBid: 3WJM shows the ***D***_3_ symmetry of this “dimer of trimers”.

## S1 Text.

**Extended details on Markov chain modelling of interaction formation and loss.**

## S2 Text.

**Parameter invariance of duplication-heteromerisation pathway and associated data collapses.**

