## Supplementary material for "Duplication accelerates the evolution of structural complexity in protein quaternary structure": S1 Text

### Markov chain modelling of interaction dynamics.

Interaction strengths can be modelled as states in a Markov process, where point mutations step between these states. With only a few assumptions, an analytic form for the baseline evolution of strength is easily achievable. The key assumptions are infinite population sizes and point mutations, which are both reasonably satisfied with the chosen parameters.

Interaction formation and loss is modelled by an absorbing Markov chain, shown with a diagram in Fig A. Interactions are formed when two initial binary strings exist below the strength threshold, and then eventually mutate above it. Transition probabilities to move up or down are directly related to the strength state and complementary weakness as $W_i = i/L$ and $W_i^\dagger = 1 - i/L$ respectively. The transition matrix $\mathbf{F}$ thus only has non-zero elements in the super- and subdiagonals, where the transition probabilities are the fractions of aligned bits (superdiagonal) and non-aligned bits (subdiagonal). The critical strength state is given by $c = \left\lceil L \cdot \hat{S}_c \right\rceil$.

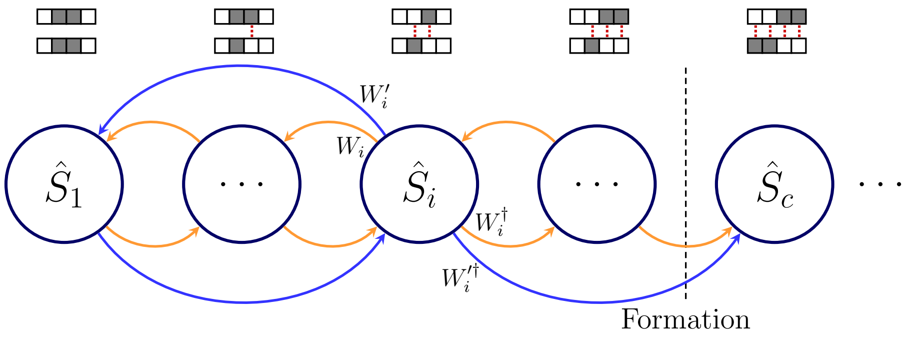


**Fig A. Example absorbing Markov chain for modelling interaction formation.** Interaction dynamics are modelled by a Markov chain of strength states, with transition weights based on interaction strength. However, now interactions are formed or lost when they walk past the strength threshold and are "absorbed". Symmetric homomeric (blue) interactions behave similarly to heteromeric interactions, albeit larger steps.

Mutations to a symmetric homomeric interaction move by two strength states on the chain per step, as any point mutation affects the actual site and its partner counter-aligned site. Equivalently, symmetric homomeric interactions have an effective binding site length of $L' = L/2$, or half as many chain states. Otherwise their interaction strength dynamics are indistinguishable from heteromeric interactions.

#### Interaction formation

The transition matrix $\mathbf{F}$ for the Markov chain model has has $N = \left\lceil L \cdot \hat{S}_c \right\rceil + 1$ terms. Once an individual first drifts above the strength threshold and forms an interaction, selection dominates and there is no drifting back below the threshold, i.e. it is absorbed, indicated by the emphasised 0 transition weight.

$$\mathbf{F} = \underbrace{\begin{pmatrix} 0 & W_1 & & & \\ W_0^\dagger & 0 & \ddots & & \\ & \ddots & \ddots & W_{c-1} & \\ & & \ddots & \ddots & \mathbf{0} \\ & & & W_{c-1}^\dagger & 0 \end{pmatrix}}_{N}$$

The properties of absorbing Markov chains can be calculated using the fundamental matrix $\mathbf{N}$, which captures the probability of transitioning between states $i$ and $j$ after exactly $k$ steps, summed over all $k$. Using the result for the infinite sum of a geometric series, the fundamental matrix can be expressed as $\mathbf{N} = (\mathbf{I} - \mathbf{F})^{-1}$, where $\mathbf{I}$ is an identity matrix. The number of expected steps to reach an absorbing state from any initial state is given by $\underline{t} = \mathbf{N}\underline{1}^\intercal$, where $\underline{1}$ is a vector of ones.

The probability to be in any given state is given by a binomial, and so the initial condition strength vector $\underline{B}$ is defined as $B_i = \binom{L}{i}(.5)^L$. The expected forming time is thus:

$$\langle \tau_{\text{form}} \rangle = \underline{B} \cdot \underline{t} = \underline{B} \cdot (\mathbf{I} - \mathbf{F})^{-1} \underline{1}^\intercal$$

Although the transition matrix appears simple, there is no general closed-form solution to the equation above. However, as suggested in the main text, the approximate empirical form derived over a realistic parameter window is:

$$\langle \tau_{\text{form}} \rangle \approx \exp\left(\left(0.7L^{-1/2} + 0.08\right) \cdot L^{1.35\hat{S}_c}\right)$$

Regardless of the exact form, there is strong dependence on both $L$ and $\hat{S}_c$ which appears quasi-exponential. Formation times quickly explode as these two parameters increase. Since symmetric homomeric interactions have an effective length of $L/2$, they routinely form significantly quicker than a heteromeric interaction with equivalent parameters.

#### Interaction loss

Due to the simplicity of binary string binding sites, interaction loss can be modelled analogously to interaction formation. Completely reversing the Markov chain and modifying which state is the critical state produces a different transition matrix that is used identically to above. Now the absorbing state corresponds to when the interaction is lost and is assumed to never reform.

Interaction strength Markov chains can be "reversed" with the anti-diagonal identity matrix (*exchange*) matrix, $\mathbf{J}$, as $\mathbf{F}^{\text{rev}} = \mathbf{J}^{-1}\mathbf{F}\mathbf{J}$. Regardless of reversing the chain, the number of states now relates to how many strengths are above the threshold, unlike in formation. The matrix then takes the size of $N = \left\lfloor L \cdot \left(1 - \hat{S}_c\right)\right\rfloor + 1$.

Initial population states are not longer binomially distributed, but are driven by selection dynamics. The steady-state distribution is given by $\underline{\pi}_{\text{PF}}$, which is the eigenvector corresponding to the largest eigenvalue of the matrix:

$$\mathbf{Q} = \underbrace{\begin{pmatrix} 0 & W_{c+1} & & \\ W_c^\dagger & 0 & \ddots & \\ & \ddots & \ddots & W_L \\ & & W_{L-1}^\dagger & 0 \end{pmatrix}}_{N}$$

See S2 Text of the generalised model's introduction [1] for more details of this derivation. The expected time to first drift below the strength threshold, and thus lose an interaction, is then:

$$\langle \tau_{\text{loss}} \rangle = \underline{\pi}_{\text{PF}} \cdot \underline{t}' = \underline{\pi}_{\text{PF}} \cdot (\mathbf{I} - \mathbf{F}^{\text{rev}})^{-1}\, \underline{1}^{\intercal}$$

While the formalisms to calculate the expected formation and loss times share mathematical similarity, the resulting values differ greatly. For all relevant parameters used in this chapter, the expected time of formation is magnitudes greater than that of loss. Additionally, while heteromeric interactions are slower to form, they are also slower to be lost. This is despite the steady-state strength distribution for heteromeric interactions skewing closer to $\hat{S}_c$ ($\langle \hat{S} \rangle_L < \langle \hat{S} \rangle_{\text{L}/2}$).
