## Supplementary material for "Duplication accelerates the evolution of structural complexity in protein quaternary structure": S2 Text

### Universality of duplication-heteromerisation pathway

With the nontrivial dependence on $L$ and $\hat{S}_c$, different model parameters can have similar expectations for one interaction type but not the other. It is critical to analyse them separately, with the first formed interaction being overwhelmingly symmetric homomeric and the second heteromeric. Rescaling the required generations to form an interaction by these expectations provides a rough measure of "evolutionary epochs", which can be more directly compared across parameterisations.

Even after collating many simulations, the distribution of formation times has massive variance and makes histogram binning difficult. Since interaction formation is essentially a Bernoulli trial, the number of attempts before it forms can be approximated with a geometric distribution. As interaction formation typically takes many generations, this geometric distribution can be replaced with its continuous version: the exponential distribution. Fitting an exponential to the data agrees well with the noisy binned data, and so is a good approximate representation that simplifies interpretation. These exponentials are shown in Fig A for unscaled and rescaled generations.

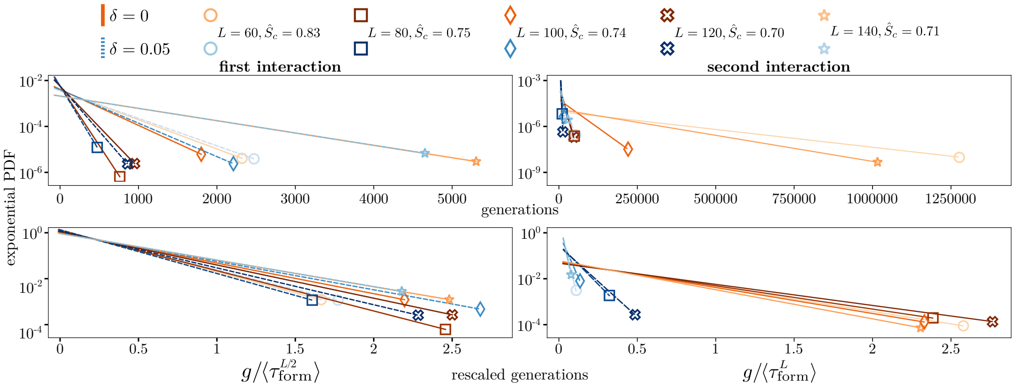


**Fig A. Interaction formation times for the first and second interactions.** Fitted exponentials approximate the distribution of formation times well for various model parameters both without and with duplication, $\delta = 0$ and $\delta = 0.05$ respectively. Unscaled generations (top) can span many orders of magnitudes for the first (left) and second (right) interactions. After rescaling by the symmetric homomeric or heteromeric expectation, the majority of simulations collapse. However, the second interaction with duplication does not collapse, and is steeper than their without duplication counterparts—equivalent to faster formation.

As predicted earlier, duplication only accelerates evolution through the heteromerisation of symmetric homomeric interactions and no gain is observed in forming the first interaction. Once rescaled, all the different parameters approximately collapse onto a single line, substantiating the theoretical expectations and providing a general principle independent of parameter choice. Similarly for the second interaction, there is a collapse of simulations without duplication. However, simulations with duplication do not collapse at all, and take significantly less time to form, showing the evolutionary acceleration gained.

As discussed in the main text, the evolutionary advantage of duplication partially arises from bypassing the heteromeric bottlneck. This bottleneck size, $\lambda$, is the ratio of formation times for a symmetric homomeric and heteromeric interaction. There is no closed-form solution for formation times, but the ratio of the empirical form gives:

$$\lambda = \frac{\langle \tau_{\text{form}}^{L} \rangle}{\langle \tau_{\text{form}}^{L/2} \rangle} \approx \exp\left(0.2L^{\left(1.35\hat{S}_c - 0.5\right)}\right)$$

As $L$ or $\hat{S}_c$ increase, the duplication pathway bypassing the bottleneck offers greater and greater gain. However, only a fraction of duplication events survive the genetic drift into the correct evolvable heteromeric state. This corresponds to when both symmetric homomeric interactions are lost but the heteromeric copy survives. Markov chains could be used again, with three correlated random walks: two of length $L/2$ and one of length $L$. An analytical result using Markov chains is difficult to derive, so these survival rates are most easily found empirically through separate simulations as $\epsilon$.

The product of these two terms, $\lambda \cdot \epsilon$, is an estimate of the evolutionary advantage of the duplication-heteromerisation pathway, and values are provided in Table 1. The three example parameterisations in Fig 6 are in complete agreement, with increasing $\lambda \cdot \epsilon$ matching an increase in $\Delta\langle\mathcal{C}\rangle$. The colour gradients in Fig A are determined by this product, with lighter colours corresponding to larger products and thus more advantage. Fitted exponential ordering reflects this, with lighter colours being steeper and thus forming the next interaction and evolving sooner.

**Table 1. Empirical duplication-heteromerisation pathway advantage.** The bottleneck size $\lambda$ varies significantly for different parameters of $L$ and $\hat{S}_c$. Similarly, the heteromeric drift survival rate $\epsilon$ depends on both parameters, but it primarily increases with decreasing $\hat{S}_c$.

| | Simulation parameters | | | | |
|---|---|---|---|---|---|
| $L$ | 60 | 80 | 100 | 120 | 140 |
| $\hat{S}_c$ | 0.83 | 0.75 | 0.74 | 0.70 | 0.71 |
| $\lambda$-ratio | 1923 | 246 | 537 | 184 | 1023 |
| $\epsilon$-factor | 0.017 | 0.033 | 0.032 | 0.045 | 0.037 |
| $\lambda \cdot \epsilon$ | 32.7 | 8.1 | 17.2 | 8.3 | 37.9 |

Since the second interaction to form when duplication is allowed is typically symmetric homomeric, it is more appropriate to rescale these times by symmetric homomeric expectations. Expectations should be weighted by the survival rate $\epsilon$, as not all duplication-heteromerisation attempts survive. The "fully" rescaled exponential fits are shown in Fig A for the second formed interactions.

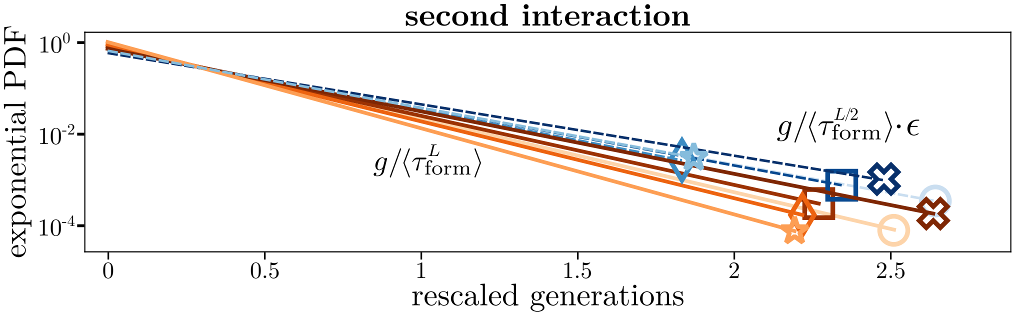


**Fig A. Fully rescaled fitted distributions and associated collapse.** Rescaling the second interaction in the duplication simulations by the symmetric homomeric expectation and an empirical pathway survival factor $\epsilon$ collapses the distributions onto the previous collapse when for when duplication is not allowed.

Now all simulations collapse onto a single general line, regardless of binding site or interaction strength parameters or if duplication is allowed. While the theoretical expectations and empirical factors do not perfectly match simulations, the proposed pathway and underlying mechanisms reasonably explain all observations in the simulations. As such, it is clear that the evolutionary advantage gained through duplication in these simulations originates from this specific proposed pathway.
